# Water soluble, red emitting, carbon nanoparticles stimulate 3D cell invasion via clathrin-mediated endocytic uptake

**DOI:** 10.1101/2021.07.31.454585

**Authors:** Udisha Singh, Aditya Guduru Teja, Shanka Walia, Payal Vaswani, Sameer Dalvi, Dhiraj Bhatia

## Abstract

Bright, fluorescent nanoparticles with excitation and emission towards the red end of the spectrum are highly desirable in the field of bioimaging. We present here a new class of organic carbon-based nanoparticles (CNPs) with robust quantum yield and fluorescence towards the red region of the spectrum. Using organic substrates like para-phenylenediamine (PPDA) dispersed in diphenyl ether and reflux conditions, we achieved scalable amounts of CNPs of the average size of 25 nm. These CNPs were readily uptaken by different mammalian cells, and we show that they prefer clathrin-mediated endocytosis for their cellular entry route. Not only can these CNPs be specifically uptaken in cells, but they also stimulate cellular processes like cell invasion from 3D spheroid models. These new class of CNPs, which have sizes similar to proteinaceous ligands, hold immense potential for their surface functionalization, whereby they could be explored as promising bioimaging agents for biomedical imaging and intracellular drug delivery.

## 1. Introduction

Bioimaging of specific biomolecules or cellular processes or cell entire cells or tissues is one of the best non-invasive ways to understand and visualize biological activity at the cellular level^1^. Recent years have witnessed the emergence of a plethora of fluorescent nanomaterials, having gained much attention in the field of bioimaging owing to their small sizes, low cost, high tuneable fluorescent properties, and their ease of interface with biological systems leading to widespread applications.^2,3^ Many different fluorescent nanomaterials have already been used, such as rare-earth semiconductor quantum dots^4,5^, organic dyes^6,^ and polymer dots^7^.

However, their applications are still extremely limited due to their limited stability, toxicity, poor water solubility, thus hindering their use for biomedical and biological applications. Carbon-based nanomaterials, especially carbon dots (CDs, including carbon nanoparticles, graphene quantum dots, and carbon quantum dots), have gained much attention because of their small size, biocompatibility, low toxicity, and stable fluorescent properties, and low cost of synthesis.^8,9^ All these carbon nanomaterials have sp^2^ hybridized carbon core and surface functional groups.^10^

Much research had already been done in the synthesis method, doping and functionalization, tuneable emission, etc. in CNPs. Most of CNPs synthesised are usually showing fluorescence in the blue-green region, which are not of much use in bioimaging because their excitation being in the UV region, which not only induces autofluorescence to disturb the signal of CNPs but also damages the cells and tissues.^11,12^ Very few reports of carbon nanoparticles showing fluorescence in the near-infrared region has been reported. For example, Hui et al. have synthesised red CDs with optical emission at 627 nm.^13^ Xiong and co-workers have synthesised red emissive CDs with optical emission at 625 nm by PPDA and urea using the hydrothermal synthesis method.^14^ Although few red emissive CNPs have been obtained showing fluorescence in the near-infrared region, the product obtained are in tiny amounts, and the process of synthesis can be tedious, limiting the use of these CNPs.

We modified the existing methods in this direction by using para-phenylenediamine (PPDA) as precursor material to achieve the synthesis of red emissive fluorescent CNPs. The CNPS obtained by reducing of PPDA exhibit excitation and emission spectra in the red region of visible spectrum. The CNPs synthesised by one step reflux reaction in a round bottom flask, which is simple, cost-effective and a large amount of product is obtained. These CNPs were seen to be readily uptaken by different mammalian cells via a very specific endocytic route called clathrin-mediated endocytosis (CME). The long-wavelength emission helps in avoiding the strong absorption from biological samples as well as prevent enhanced emission due to autofluorescence from the cells. Not only were the CNPs actively uptaken by cells, but they also stimulated different cellular physiological processes like cell invasion in 3D spheroid models. Overall, our results indicate that the class of CNPs presented here are good bioimaging material and can lay a path in the development of advanced bioimaging tools.

**Scheme1:**
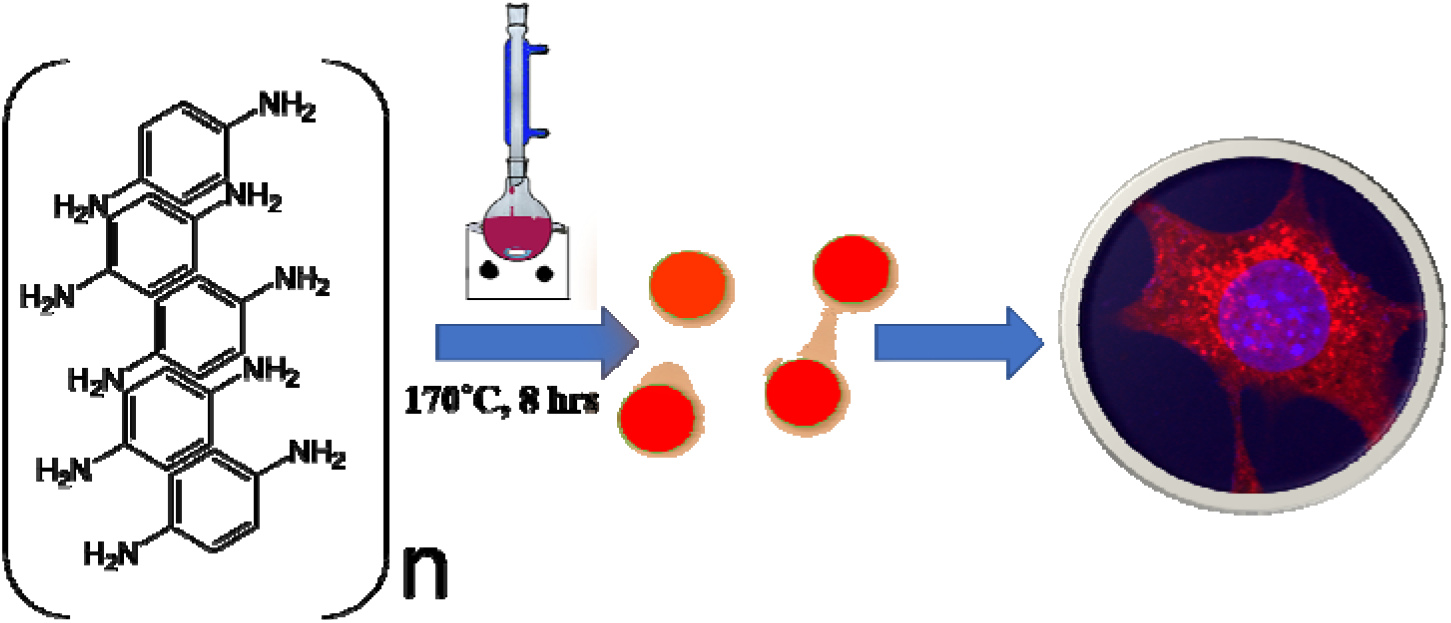
Schematic representation of CNPs synthesis using para-phenylenediamine dissolved in diphenyl ether, refluxed at 170□C for 8 h leading to one-step formation of red emitting nanoparticles that could be uptaken by cells for bioimaging applications.

## 2. Results and Discussion

### 2.1. Synthesis and Characterization of CNPs

Red fluorescence emitting CNPs were synthesised through reflux reaction mediated pyrolysis of PPDA in diphenyl ether for 8 h at 170°C. The size and morphology of the CNPs were characterized using TEM. TEM images displayed that these particles are adequately dispersed and possess a spherical disk-like structure. The mean diameter of the fluorescent CNPs is 26.7 ± 3.256 nm while counting 100 nanoparticles displayed in **Figure 1a**. The particle size measurement was done by Image J software using a scaled image and analysed using Origin software. Through the HRTEM image in **Figure 1b,c** lattice fringes with a lattice distance of 0.2075 nm, which is in good accordance with the (101) planes of carbon were obtained suggesting that CNPs synthesised are crystalline in nature.^16^ AFM image discloses the topological heights of 2.6 nm, indicating that fluorescent CNPs contain an average of 7 to 8 layers of graphene sheets (**Figure 1d**).^17^

**Figure 1:**
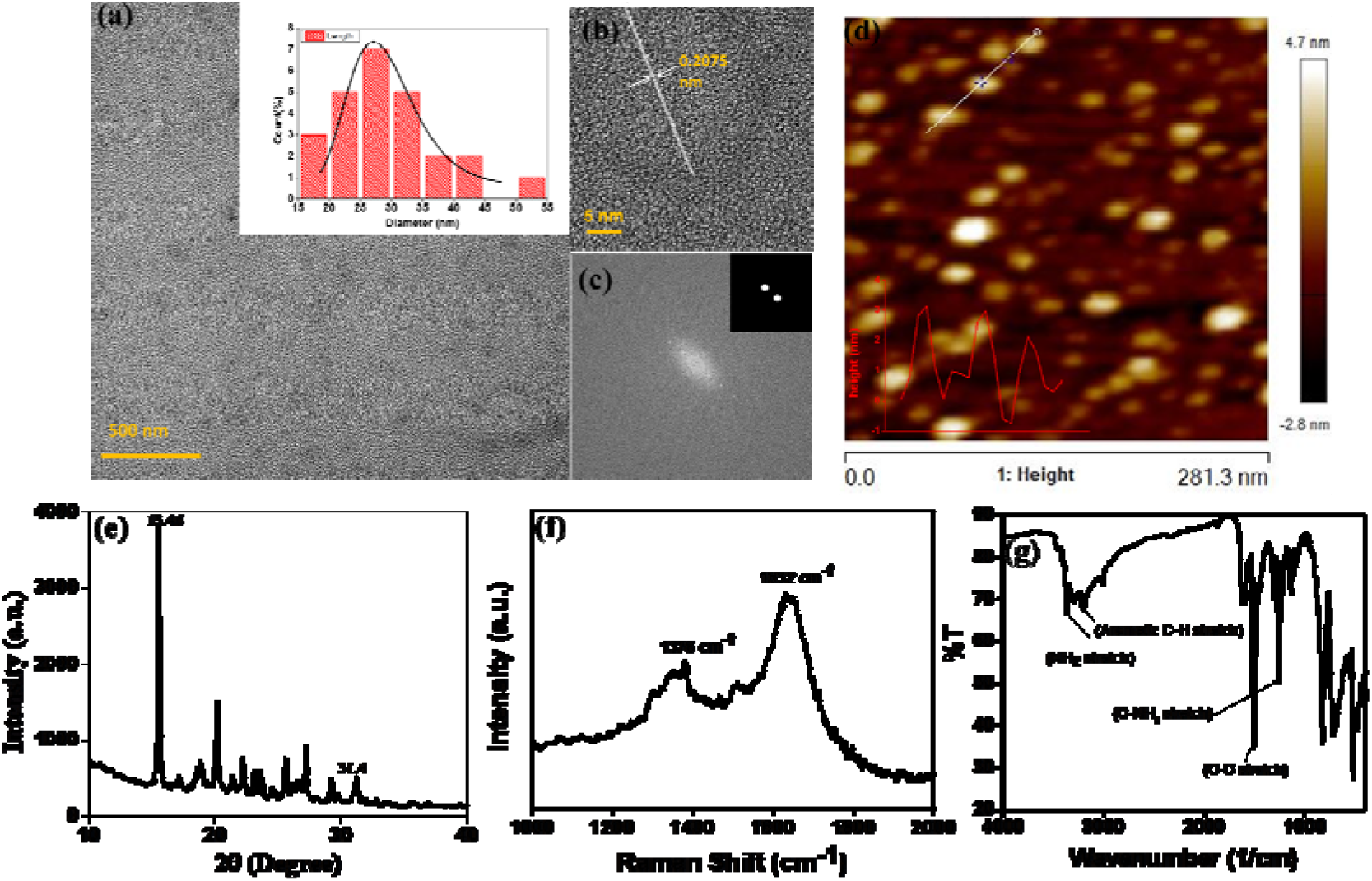
Characterization of fluorescent CNPs. **(a)** TEM image shows spherical-disk shaped CNPs of size 26.7 ± 3.2 nm **(b)** HRTEM image of CNPs showing lattice fringes of 0.2075 nm **(c)** Fast Fourier Transform image of CNPs exhibiting (1 0 1) lattice planes **(d)** AFM image of CNPs showing the similar sized CNPs with the topological heights of 2.6 nm **(e)** XRD spectra of CNPs showing that CNPs are crystalline **(f)** Raman spectra of CNPs giving D and G band value of 1378 cm^-1^ and 1632 cm^-1^ respectively **(g)** FTIR spectra of CNPs showing the functional groups presenting on the surface of CNPs

To further understand the structure and composition of CNPs, we performed XRD, Raman and FT-IR spectroscopy of fluorescent CNPs. The XRD profile of CNPs shows several peaks that may be due to the ordered stacking of synthesised CNPs. The number of peaks also shows the complex mixture of crystalline components in the sample solution of CNPs.^18^ The peaks at 15.46° and 31.4° correspond to the precursor material PPDA, revealing some impurities are present due to incomplete carbonization of PPDA as shown in **Figure 1e**.^19^ Raman spectroscopy was also used as a non-destructive way to understand the degree of graphitization of CNPs. We obtained two peaks at 1378 cm^-1^ and 1632 cm^-1^ representing the D band and G band respectively (**Figure 1f**). The D band constitutes the vibrations of disordered graphite or glassy carbon. The G band represents the in-plane displacement of carbon atoms in a two-dimensional hexagonal lattice. Also, the value of I_D_/I_G_ is being used to evaluate the degree of graphitization.^20^ The intensity of the G band is higher than the peak of the D band with the I_D_/I_G_ value of 0.844 (**Figure 1f**), implying that more sp^2^-hybridized carbon present as compared to sp^3^ hybridized carbon. FTIR spectrum gives a qualitative idea about the functional group present on the surface of CNPs. CNPs mainly contain C, O and N elements forming chemical bonds -NH_2_ (3364 cm^-1^), C-C (1510 cm^-1^) and C-NH_2_ (1257 cm^-1^) as shown in **Figure 1g**. The absorption peak obtained at 3364 cm^-1^ confirms the presence of the amine group on the surface of FCNPs, also proving the nitrogen doping of CNPs. The C-H stretching occurs around 3180 cm, ensuring that the CDs contain the aromatic ring of carbon.

### 2.2 Optical properties of CNPs reveal stable, pH responsive, red emitting fluorescence

To understand the optical properties of CNPs, UV-visible absorption and fluorescence emission, and excitation spectra were measured. Through UV-vis spectra, we obtain three absorbance peaks at 241 nm, 306 nm, and 415 nm. (**Figure 2a**) The absorbance peak at 241 nm assigned to π-π* transition from aromatic carbon structure, the shoulder peak at 306 nm assigned to π-n* transition from functional groups with lone pairs electron and the absorbance peak at longer wavelength 415 nm assigned to the energy level transition from the angstrom-sized conjugated π-structure present in CNPs.^21^ The fluorescent emission spectra of fluorescent CNPs was measured as a function of different excitation wavelength. The fluorescence (FL) spectra of CNPs gives a maximum peak at 622 nm with an excitation wavelength of 480 nm. As shown in (**Figure 2b**), the FL spectra of CNPs don’t shift with respect to change in excitation wavelength of 400 to 520 nm range, indicating that fluorescent CNPs have distinct excitation independent FL behaviour, which is very seldom observed. This also shows that the CNPs exhibits a quiet homogenous size distribution, composition and surface state. The sp^2^ hybridized carbon network in CNPs can also be one of the reasons for independent excitation fluorescence. The CNPs are shown brownish-red colour under white light and shows the dark red colour under UV light illumination in aqueous solution, as shown in the inset of **figure 2a**.

**Figure 2.**
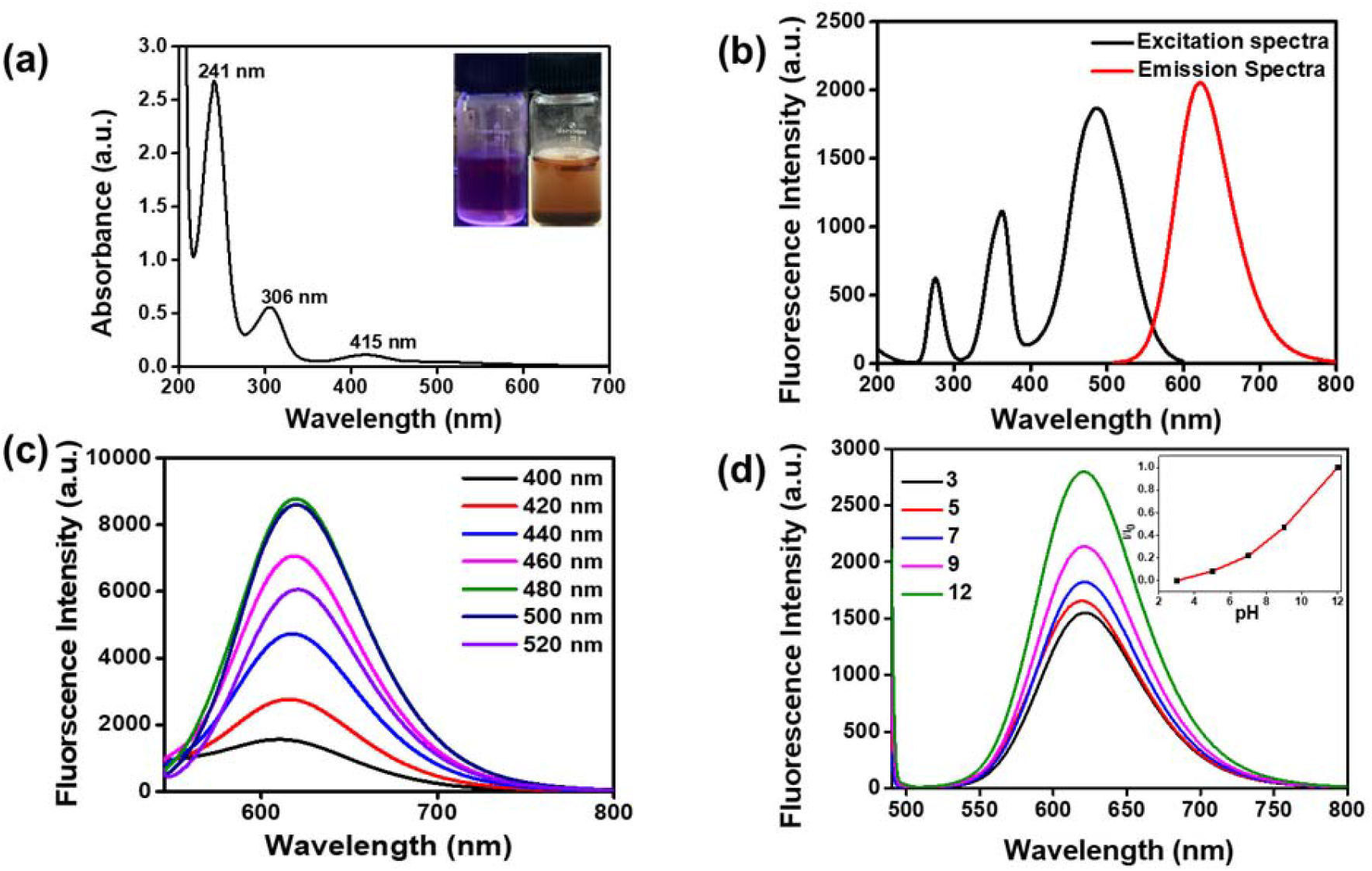
**(a)** UV-Visible spectra of CNPs showing three absorbance peak-241 nm, 306 nm and 415 nm π-π* transition from aromatic carbon structure, π-n* transition from functional groups with lone pairs electron and the last one assigned to the energy level transition from the angstrom-sized conjugated π-structure present in CNPs **(b)** Excitation and emission spectra of CNPs, at 480 nm excitation wavelength showed λ_max_ at 622 nm wavelength **(c)** Emission spectra CNPs at different excitation wavelength ranging from 400 nm - 520 nm with emission at λ_max_ of 622nm with varying intensities **(d)** Emission spectra of CNPs dispersed in different pH solutions (3,5,7 and 12).

The fluorescence quantum yield of optimal CNPs comes out 5.86% in milli-Q water and 87% in ethanol by referring to standard rhodamine B (QY=33%). This also shows that the synthesised CDs are more dispersed in organic solvents and shows higher QY and fluorescence. The CNPs synthesised were redispersed in different solvents-water, hexane, acetone, acetonitrile, DMSO, DMF, ethyl acetate, IPA, ethanol and methanol. All these ten solvents have different polarities, and their emission spectra also shift with a change in solvent, as shown in **supplementary information Figure S1a**. The colour of the CNPs changes in different solvents showing that the fluorescence of CNPs not only dependent on functional groups present on their surface but also on the type of dispersant used.^15^

For cellular stability applications, the pH-dependent studies of CNPs were done by monitoring change in their fluorescence signal as a function of pH. Different pH solutions in milli-Q water were prepared (3, 5, 7 & 12) and CNPs (0.5 mg/ml) were dispersed in it, and emission spectra were taken as shown in **Figure 2d**. The fluorescent CNPs shows an increase in fluorescence with an increase in pH, hence showing great potential in pH sensing. The lowest fluorescence intensity was found at pH 3 and the highest intensity found at pH 12. The percentage increase in intensity from pH 3 to pH 7 is 38% whereas the increase in fluorescence intensity from pH 7 to pH 12 is approximately 31%. The pH sensitivity of CNPs originating may be due to protonation and deprotonation of oxygen-containing groups on the surface.^22^ Also, there is a significant colorimetric change of CNPs dispersed in different pH solutions.

For application in cellular bioimaging, the stability of fluorescent CNPs in different ionic concentrations is very crucial. To check the fluorescence stability of CNPs under a highly ionic environment, we dispersed CNPs (0.5 mg/ml) in different concentrations of KCl salt (0 to 1 M) solution. Our results show a significant increase in fluorescence of CNPs with an increase in the concentration of KCl (**Figure 3a**). We also analysed the photostability of CNPs under constant visible light at 480 nm. The CNPs were irradiated with uninterrupted excitation wavelength light for 90 mins, and readings were taken at an interval of every 10 mins. The resultant fluorescent intensity (I/I_o_) stipulates insignificant decrease in fluorescence with time, indicating the photostability of synthesized CNPs **Figure 3b**.

**Figure 3.**
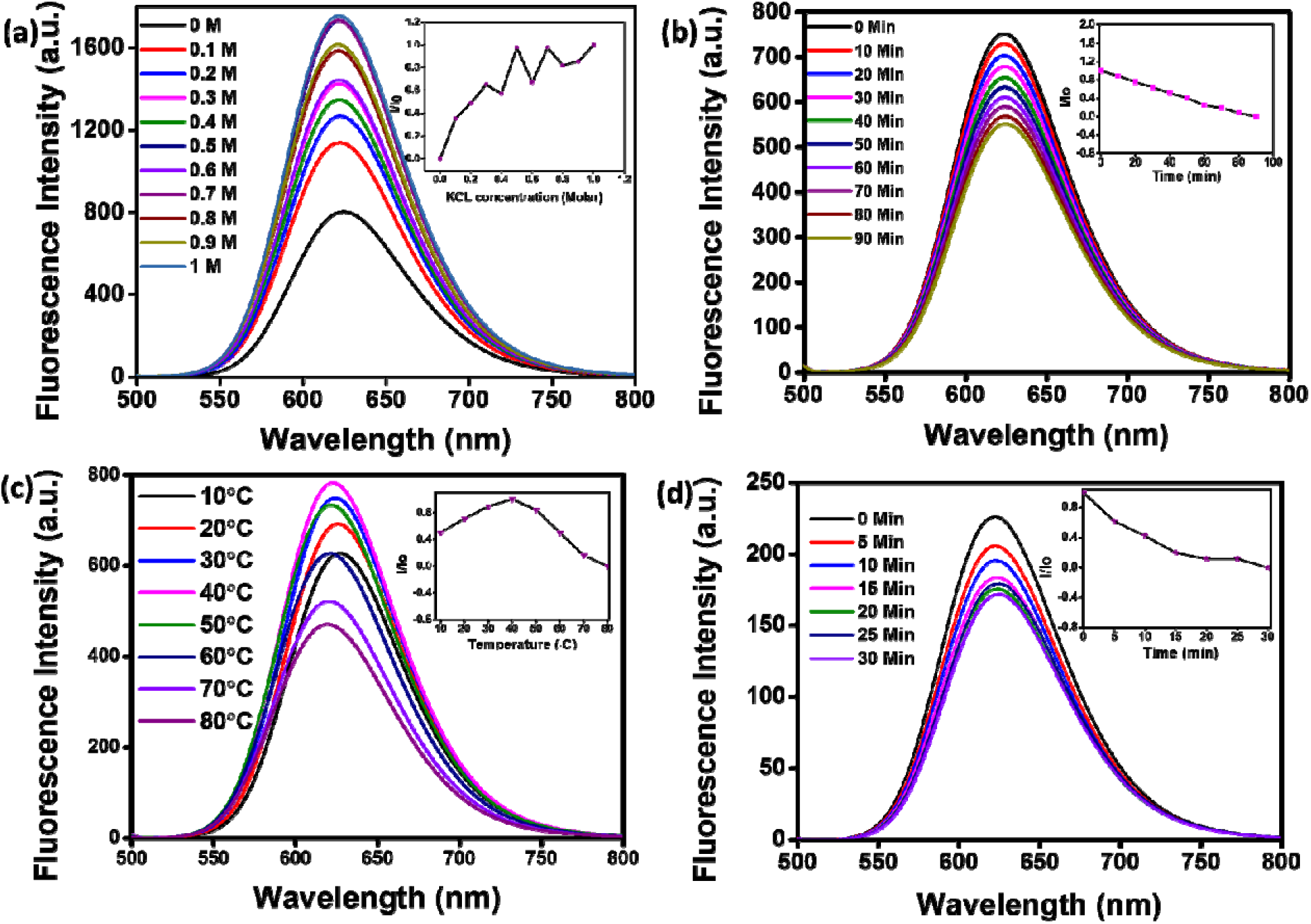
Photostability studies of CNPs. **(a**) Fluorescence emission spectra of CNPs titrated with different ionic concentrations of KCL (0 to 1M) **(b)** Fluorescence spectra of CNPs recorded from 0 to 90 min at the increment of 10 min, irradiated with an excitation wavelength of 480 nm continuously to check their photostability. **(c)** Fluorescence spectra of CNPs at different temperature range (10°C-80°C) recorded at an increment of 10 °C. **(d)** Fluorescence spectra of CNPs at different time points (0-30 min) when kept at 80°C. Inset showing relative effect on the photostability of CNPs excited under various condions

We further studied the thermal stability of CNPs using a fluorescence spectrophotometer and change the temperature from 10°C to 80°C. With increase in temperature, initially the fluorescence intensity increased from 10°C to 40°C followed by decrease from 50°C to 80°C (**Figure 3c**). The increase in fluorescence intensity from 10°C to 40°C is 20%, and the decrease in intensity from 50°C to 80°C is 40%, indicating that the fluorescence intensity of CNPs is temperature-dependent and the best fluorescence is obtained at physiological temperatures of cells and tissues. The CNPs were also kept at 80°C for 30 minutes to check their thermal stability. There was an insignificant decrease in fluorescence intensity of CNPs with time at 80°C (**Figure 3d**). The reduction in fluorescence intensity with an increase in temperature from 40°C to 80°C can be explained by thermally activated non-radiative channels. At high temperatures, the non-radiative channels are activated, leading to exciting electrons not effectively releasing photons. That is why we see a reduction in PL intensity at high temperatures.^23^

### 2.3. Cellular uptake of CNPs follow clathrin-mediated endocytosis

To check the suitability of the developed CDs as molecular imaging probes toward biological systems and also to monitor the effect of the same on the normal and cancer cells, MTT assay was performed with SUM159A and MEF cells (**supplementary Figure S2 a,b**). The experiments were carried out using different concentrations, namely 10, 20, 30, 50, 100, 200, and 500 µg/ml of CNPs. After 24 h of incubation with CNPs, SUM159A showed ∼80% cell viability up to 50 µg/ml concentration, and then decreased to ∼62% up to 200 µg/ml and finally up to 52% at 500 µg/ml of CNPs concentration (**supplementary Figure SI 2a**). In the case of MEF, the CNPs showed cell viability of 70% and 50 % at a concentration of 10-20 µg/ml and 30-100 µg/ml, respectively. The cell viability percentage significantly dropped, documented ∼37% and 16% cell viability at 200 and 500 µg/ml concentration of CNPs. The cytocompatibility studies with SUM159A suggested that CNPs did not show any toxicity up to 100 μg/mL even after 24 h, whereas the same limit for MEF was found to be 20 μg/mL. This also suggests that different cells have different capacities to interact with CNPs, thus affecting their cytotoxicity.

To explore the cellular uptake properties of CNPs, confocal microscopy studies were done to analyse the time and concentration-dependent cellular internalization of CNPs using SUM159A and MEF cells (**Figure 5 and Figure SI 3i, respectively**). Cells, when incubated with CNPs of different concentrations at 37°C and analysed by confocal microscope, revealed the successful internalization of CNPs. Further, confocal images along with quantification data suggested that CNPs showed strong intracellular fluorescence response with respect to time and concentration. Concentration-dependent studies showed that the fluorescence intensity was increased ∼1-2 times from 10 to 100 µg/ml concentration of CNPs. The cell morphology was retained unchanged at lower concentrations. However, when cells were exposed to a higher concentration of 100µg/ml of CNPs, the fluorescence signal was observed from the nucleus, and cells appeared in stress. (**Figure 5a and supplementary Figure SI 3i**). Further, time-dependent studies showed an increase in fluorescence signal up to 60 min with enhanced fluorescence intensity (**Figure 6 and supplementary Figure SI 3ii**). Confocal microscopic studies showed that the synthesized CNPs could be used for bioimaging applications due to their strong and stable fluorescence emission in the red channel.

**Figure 5:**
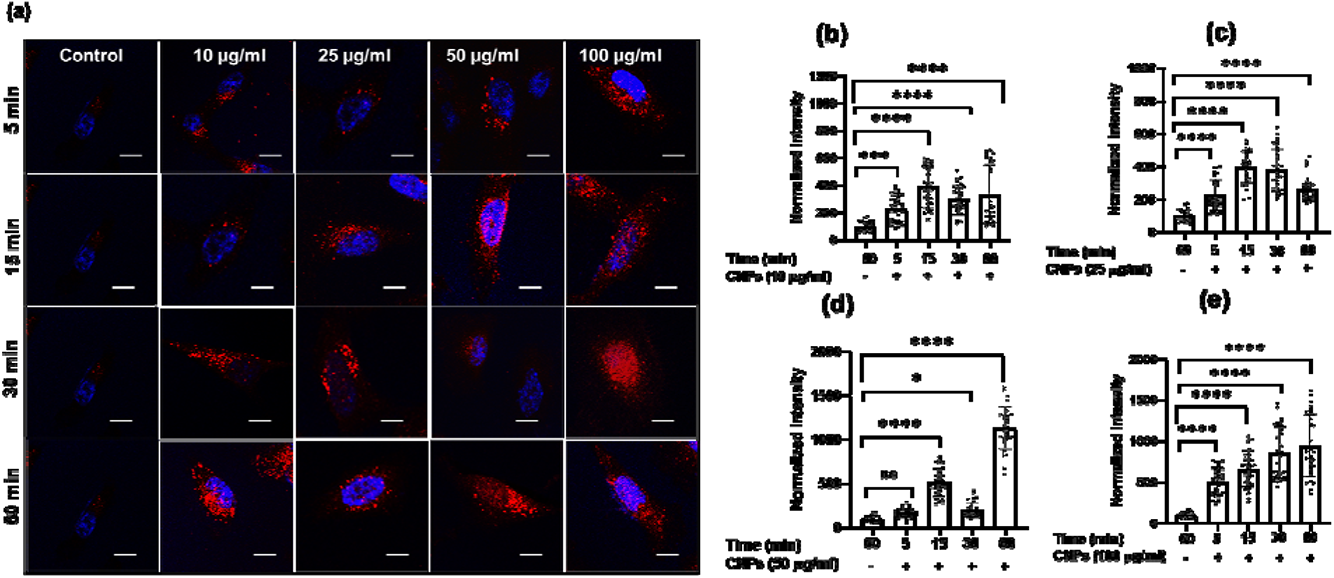
Concentration-dependent cellular uptake studies of CNPs at different time intervals (a-d) Confocal images of SUM159A incubated with CNPs for 5, 15, 30, and 60 min, respectively. Scale bar is 10 µm for all the images. The first column in the figure represents untreated SUM-159A cells imaged after 60 min of incubation. (e-h) Quantification of cellular uptake of CNPs at 5, 15, 30, and 60 min, respectively. The scale bar is 5 µm. **** Indicates statistically significant value of p < 0.0001, *** Indicates statistically significant value of p = 0.0006, ** Indicates statistically significant value of p = 0.0074 and * = 0.03 (one-way ordinary ANOVA).

**Figure 6.**
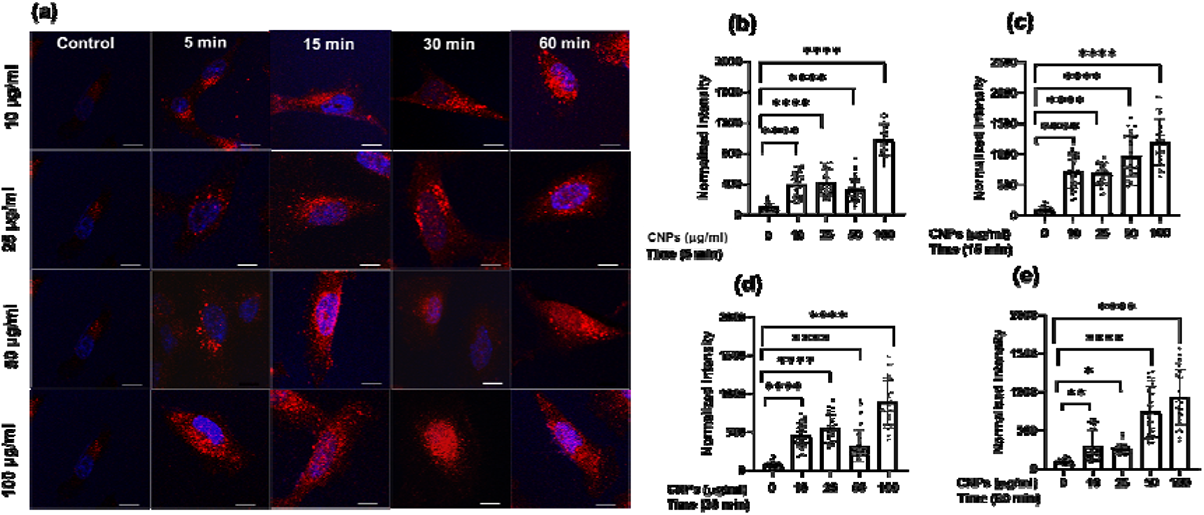
Time-dependent cellular uptake studies of CNPs at different time intervals. (a) Confocal images of SUM 159 incubated with 10, 25, 50, and 100 µg/ml of CNPs, respectively. The first column in the figure represents untreated SUM-159 cells imaged after 60 min of incubation. Scale bar is 10 µm for all the images. (b-e) Quantification of cellular uptake of CNPs at 10,25, 50, & 100 µg/ml, respectively. The scale bar is 5 µm. **** Indicates statistically significant value of p < 0.0001, ** indicates statistically significant value of p = 0.002. and * = 0.01 (one-way ordinary ANOVA).

Clathrin-mediated endocytosis (CME) and clathrin-independent endocytosis (CIE) are the most important pathways considered for NPs uptake.^24^ We thus investigated the plausible endocytosis pathway responsible for the cellular uptake of CNPs by using cellular cargoes markers viz., Transferrin (Tf), a well-known target of CME, whereas Galectin3 (Gal3) for marking CIE were included as a reference. Specific inhibitors, namely dynasore to block CME and lactose for CIE inhibition, were used to validate their effect on CNPs uptake. When cells were pulsed with these markers and CNPs in the presence of these inhibitors, we observed that Tf and CNPs uptake was significantly reduced by dyansore treatment in SUM-159A cells (**Figure 7a, b**). Whereas lactose inhibition resulted in a non-significant change in fluorescence intensity in the case of Tf as well as CNPs, while the intensity of Gal3 uptake was significantly decreased.

**Figure 7:**
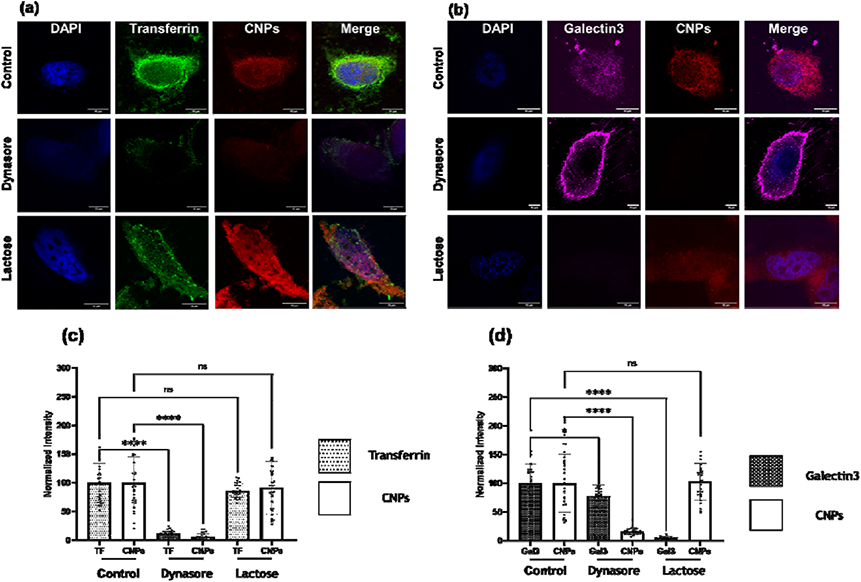
Uptake of CNPs via endocytosis pathway in SUM-159 cells under conditions that perturb CME and CIE endocytosis. (**a**) Uptake of CNPs and A488-labelled transferrin (Tf) in cells treated with dynasore (80CµM), and lactose (150 mM). (b) Quantification of normalized intensity. (c) Uptake of CNPs and A647-labelled galectin-3 (Gal3) in cells treated with dynasore (80CµM). and lactose (150 mM). Scale bar is 10 µm for all the images. (d) Quantification of normalized intensity. **** Indicates the statistically significant value of p < 0.0001 * Indicates statistically significant value of p = 0.376 and ns indicates non-significant value of p (one-way ordinary ANOVA

We also examined the cellular uptake of CNPs via the CIE pathway using Gal3 as a reference marker. Lactose is known to compete with extracellular Gal3 for binding to specific β-galactosides on the plasma membrane receptors and, therefore, used to inhibit the uptake of extracellular Gal3.^25^ It was observed that the uptake of Gal3 was significantly decreased when inhibited by lactose (CIE blocker), whereas CNPs uptake was marginally affected in cells (**Figure 7c, d**). However, on inhibition via dynasore, the uptake of Gal3 was reduced only marginally. On the contrary to this, CNPs uptake was decreased significantly upon treatment with dynasore, indicating the strong involvement of Dynamin which plays the role of scission of the clathrin-coated vesicles formed at the plasma membrane. These results confirmed our previous observation that CNPs uptake was CME dependent.

### 2.7. CNPs uptake stimulate cellular invasion in 3D spheroids model

Three-dimensional (3D) spheroids-based cancer models are well explored to study tumour progression, cell activity, and the effect of the pharmacokinetic drugs. Three-dimensional spheroids provide better insight into disease state, proliferation, invasion, or migration of tumour cells. Therefore, we further explored the cellular uptake of CNPs by 3D spheroids and its effect on cell invasion in a 3D matrix. Spheroids were made from triple-negative breast cancer MDA-MB-231 cells using a hanging drop method, and the well-formed spheroids were suspended in collagen with or without CNPS. The spheroids were allowed to grow in the matrix for 24h at 37°C, post which they were fixed, stained for actin cytoskeleton using Phalloidin green, and imaged under the confocal microscope. Spheroids treated with three different concentrations of CNPs viz., 50, 100, and 200 µg/ml were incubated for 24 h at 37 °C (**Figure 8a, b**). Cells, while migrating in 3D, had actively endocytosed the CNPs from the collagen matrix, probably in a CME pathway. To study the effect of CNPs on the invasion capacity of cells in 3D spheroid models, the invasion index was plotted, which showed 3D tumour models treated with 100 µg/ml of CNPs had a cell invasion index of 0.05, which was higher than the cell invasion index obtained for the tumour model for 50 µg/ml and control (**Figure 8c**). In 200 µg/ml, there was no proliferation observed, indicating an inhibitory effect of CNPs on cell invasion at higher concentrations.

**Figure 8.**
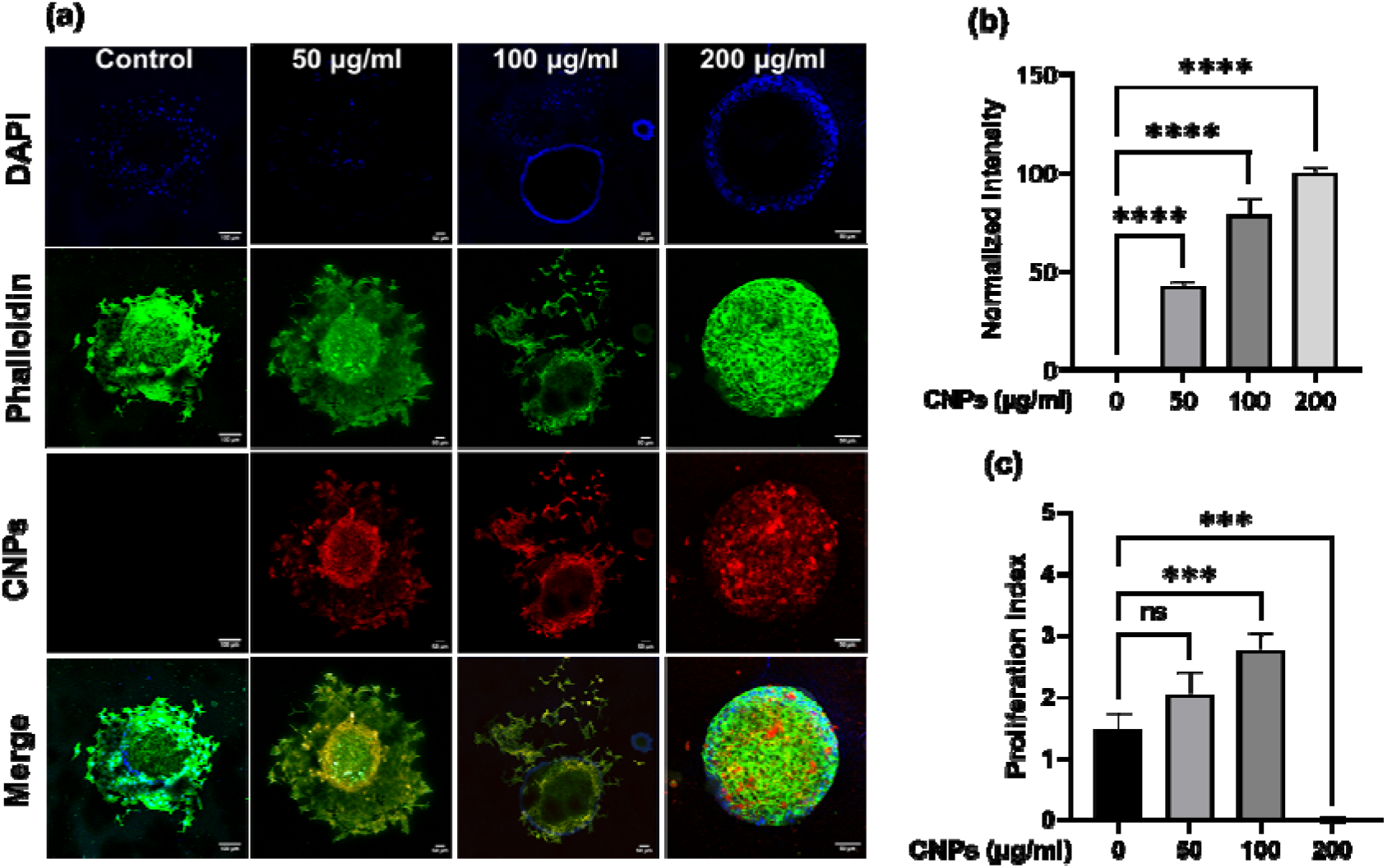
**(a)** Confocal microscopy images of 3 D spheroids treated with 50, 100, and 200 µg/ml of CNPs for 24 h The scale bar is 100 µm in the control sample and 50 µm for the rest of the images. Scale bar is 100 µm for merge images and 50 µm for all other channels. (b) Quantification data of normalized intensity of CNPs. (c) Quantification data of proliferation index. **** Indicates statistically significant value of p < 0.0001, *** Indicates statistically significant value of p < 0.001 whereas ns indicates non-significant data (one-way ordinary ANOVA). N = 3 spheroids per condition.

## 3. Conclusions

We designed near-infrared red emissive CNPs through the reflux reaction method of heating using para-phenylenediamine (PPDA) as precursor material and diphenyl ether as solvent. Owing to red emissive fluorescence and good biocompatibility, the CNPs have great potential in the field of bioimaging. Different characterization techniques were used to properly analyse the structure, morphology, and optical properties of CNPs. The QY of CNPs comes out to be ∼ 6% in water and 87% in ethanol, showing that the fluorescence of CNPs also depends on the type of solvent used. The fluorescence intensity of CNPs also depends on the pH of the dispersing medium (water), which increases with an increase in pH. Ionic stability, photostability, and thermal stability were also checked. In cell viability studies, we observed that CNPs did not show any toxicity up to 100 μg/mL even after 24 h in SUM159A cells. Concentration-dependent and time-dependent studies on cellular uptake of CNPs were done and observed that most CNPs were uptaken into the cells through clathrin-mediated endocytosis. The same effect was observed not only in 2D but also in the 3D spheroid model, and it was observed that CNPs could trigger the invasion of MDA-MB-231 cells in a collagen matrix. The present work provides not only promising fluorescent nanomaterial for bioimaging but also opens other platforms for modulating the surface properties of nanoparticles along with this capacity to bioconjugate to different biomolecules, which will have widespread biological and biomedical applications of CNPs in coming times.

## 4. Materials and methods

### 4.1 Materials

Para-phenylenediamine (PPDA), moviol, Hoechst, dynasore, Galectin3 (Gal3) (Alexa 647), Transferrin (Tf) -A488, 3-(4,5-Dimethylthiazol-2-yl)-2,5-diphenyltetrazolium bromide (MTT), were obtained from Sigma-Aldrich. Diphenyl ether (99%) obtained from Avra, n-hexane of HPLC grade, acetone (>99.5%), N,N-dimethylformamide (>99.5%), ethanol(>99.5%), methanol (>99.8%), acetone, acetonitrile, Isopropyl alcohol(IPA) and lactose were purchased from Merck. Paraformaldehyde, Dimethyl sulfoxide (DMSO), Rhodamine B, and cell culture dishes for adherent cells (treated surface) were procured from Himedia. Ham’s Nutrient Mixture F12 (HAMs F12), Dulbecco’s modified Eagle’s medium (DMEM), fetal bovine serum (FBS), penicillin−streptomycin, trypsin−EDTA (0.25%), and collagen1 rat tail were purchased from Gibco. All the chemicals were of analytical quality, and no further purification was required.

### 4.2 Synthesis of Fluorescent CNPs

Highly fluorescent red light-emitting CNPs were synthesised using the pyrolysis method by reflux reaction.^15^ We have modified the experimental conditions to obtain the CNPs of desired size and properties. Para-phenylenediamine (PPDA) is used as precursor material and dissolved in diphenyl ether. The reaction takes place for eight hours at 170°C. After the heating process, the product was allowed to cool down at room temperature. The CNPs synthesised were precipitated out using hexane by solvent displacement method. The product obtained after heating adds dropwise to 100 ml hexane solution for the separation of CNPs. The precipitated CNPs were centrifuged and washed three times using hexane and then kept in normal atmospheric condition to dry up.

### 4.3 Various analytical methods used to study the characterization of CNPs

The shape and size of the fluorescent CNPs were analysed using (FEI Titan Themis transmission electron microscope (TEM) 60-300 kV). Atomic force microscope (AFM) provides both two-dimensional (2D) and three-dimensional(3D) images of CNPs. X-ray diffraction (XRD) technique was used to characterize the crystalline nature of CNPs and for phase identification. The scans from X-ray diffraction spectroscopy (Brüker-D8 DISCOVER) were recorded with a speed of 0.2°/min from 10° to 60° with Cu-K_α_ radiation. Raman spectroscopy was measured using a (Kaiser Raman spectroscopy) with a 785 nm wavelength of YAG laser. The fluorescence spectra obtained using FP-8300 Jasco spectrofluorometer (Japan) in the excitation range of 400 to 520 nm. Rhodamine B was used as a standard reference dye for fluorescence measurement, with a relative quantum yield of 5.86% in water and 87% in ethanol. UV-VIS absorbance spectra of CNPs obtained by using Spectrocord-210 Plus analytikjena (Germany). Fourier transform infrared spectroscopy (FT-IR) of CNPs were recorded by using FTIR spectrometer from Spectrum 2, PerkinElmer in ATR mode. Scanning was done in the range of 400 cm^-1^ to 4000 cm^-1^. The cell imaging for fixed samples was performed under 63X resolution using confocal laser scanning microscopy platform Leica TCS SP8.

### 4.4 Spectroscopic studies

The solvents used to record absorption and emission spectra are of spectroscopic grade. The emission spectra were recorded 10 nm after the excitation wavelength for the standard recording of fluorescence spectra. Quantum yield were determined by comparison with rhodamine B in ethanol (□ = 0.33) as reference using the following equation:

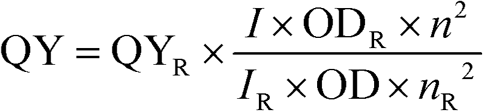

QY is the quantum yield, *I* is the integrated fluorescence intensity, *n* is the refractive index, OD is the optical density of that particular excitation wavelength, and R is used for reference.

### 4.5 Cell Culture

Mesenchymal triple-negative breast cancer cell line (SUM159A) and Embryonic Mouse Fibroblasts (MEFs) cells were obtained as a gift from Prof. Ludger Johannes, Institut Curie, Paris France. SUM159A and MEF cells were cultured in HAMs F12, and DMEM supplemented with 10% FBS and 1% penicillin-streptomycin. For all the studies, PBS of 1× strength with pH 7.4 was used.

#### 4.5.1 MTT assay

To assess the toxicity of the CNPs, the cells were seeded in a 96-well plate with a cell density of 10,000 cells per well (100 µl) and incubated for 24 h in a CO_2_ incubator, maintained at 5% CO_2_ and 95% humidity, for cell adherence. After that, the cells were washed with PBS and treated with 100 µl of 4 different concentrations of CNPs viz., 10,20,30,50,100,200, & 500 µg/ml, in triplicates and were incubated for 24 h. After the completion of incubation time, a colorimetric 3-(4,5-dimethylthiazol-2-yl)-2,5-diphenyltetrazolium bromide (MTT) assay was performed to quantify the effect of CNPs on cell viability. At the end of the treatment, cells were washed twice with PBS; then 10 μL of MTT solution (5 mg/mL) was added to each well, and the solutions were further incubated for 3 h at 37°C. After incubation, the formazan crystals formed were dissolved in 100 μL DMSO and incubated in the dark for 15 min. The intensity of the colour was then measured by the absorbance at 570 nm from the formazan crystals and was representative of the number of viable cells per well. The values thus obtained for the untreated control samples were equated to 100%, and the relative percentage values for CNPs were calculated accordingly. All experiments were performed in triplicate, and the cell viability (%) was calculated using equation 1.

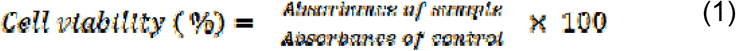

#### 4.5.2 Confocal Microscopy Studies

SUM 159 and MEF cells were seeded in HAMS F12 and DMEM at a density of ∼40,000 cells/well on coverslips in 24 well plate cell culture plates and incubated for 24 h at 37 °C and 5% CO_2_. After incubation, cells were washed twice with PBS and treated with 10, 25, 50, and 100 µg/ml of CNPs, and all the samples were further incubated for different time points of 5, 15, 30, and 60 min, respectively. After desired incubation time, the cells were washed twice with PBS and then fixed using 4% PFA for 20 min. Afterwards, cells were washed with PBS thrice, stained with DAPI, and mounted using a mounting medium (moviol). Finally, cellular internalization of CNPs was examined through Leica TCS SP5 confocal microscope with a 63× oil immersion objective. A 405 nm laser was used as the excitation source for DAPI, while a 488 nm argon laser was used as an excitation source for CNPs. The emission bandwidth for DAPI and CNPs was 420-450 nm and 590-700 nm, respectively.

#### 4.5.3 Cellular Uptake studies via endocytosis

We next wanted to explore the pathway by which CNPs were entered inside cells. Briefly, SUM-159 cells were seeded on coverslips placed in 24 well plate with a cell density of 40,000 cells/well. The cells were treated with dyansaore (80 μM) and lactose (150 mM) in HAMs-F12 (serum-free media) in desired wells and incubated for a specific time interval at 37 °C. Dyanasore and lactose were used to inhibit clathrin-mediated endocytosis (CME) and clathrin-independent endocytosis (CIE) pathways, respectively. The cells without any treatment were taken as control.

Afterwards, the cells were washed gently and incubated with fluorescently labelled markers, which follow CME (Tf(5 μg/mL and CIE (Gal3 (5 μg/mL)) pathways and CNPs (25 µg/ml) for a specific time interval at 37 °C. The cells were washed thrice with PBS and fixed using 4% PFA for 15 min at 37 °C. The coverslips were mounted on Moviol containing Hoechst for imaging. Microscopy imaging of fixed cells was performed using a confocal laser scanning microscope under 63X resolution. Quantification of data was done using Fiji ImageJ software.

#### 4.5.4 Cell invasion assay using 3D spheroid model

3D spheroids were prepared by using the hanging drop method. A T25 flask with 85 percent confluency was trypsinized with 1 mL trypsin to harvest the cells from the culture flask. Two millilitres of complete fresh medium were added to the trypsinized solution and thoroughly mixed with inactivating the trypsin. The mixture containing cells was centrifuged at 500 g for 3 minutes. The pellet obtained was resuspended in 4 mL of complete fresh medium and redispersed. Then using 35 µL of medium-contained cells were applied as drops on the inside surface of the Petri dish cover. To provide a moisture environment for spheroid formation, 15 mL of phosphate buffer saline was poured into the Petri dish. The cell droplet-containing Petri dish lid was turned upside down, covering the Petri dish, and placed for incubation at 37°C in an incubator. 3D spherical spheroids were formed after 36 h of incubation. Each drop consists of one spherical spheroid. The spheroids collected from the hanging drop are further used for cell proliferation assay. Spheroid collected from the hanging drop was added to the ECM matrix containing collagen and media in the 2:1 ratio, respectively. After the transfer to the matrix, the spheroids were subjected to the respective treatments with CNPs. Spheroid without any treatment and was subjected to complete media is considered to control. Post 24 h the spheroids were fixed using 4% PFA and stained by using Alexa flour 488 stain and Hoechst. The spheroids were mounted onto slides and further imaged under Leica confocal microscope. The proliferation index/migration index was calculated by using the formula mentioned below.

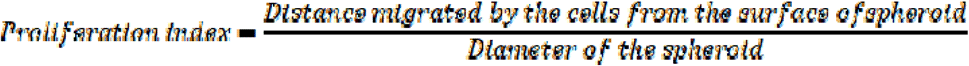

## Acknowledgments

We sincerely thank all the members of DB group for critically reading the manuscript and their valuable feedback. US, AT thank IITGN-MHRD, GoI for PhD and SW thanks IITGN-MHRD postdoctoral fellowship. PV acknowledges PhD fellowship from UGC-CSIR, India. We thank Prof. Abhay Raj Goutam for help with TEM imaging. DB thanks SERB, GoI for Ramanujan Fellowship, IITGN, for the startup grant, and DBT-EMR, Gujcost-DST, GSBTM and BRNS-BARC for research grants. Imaging facilities of CIF at IIT Gandhinagar are acknowledged.

## Supplementary Information

**SI 1:**
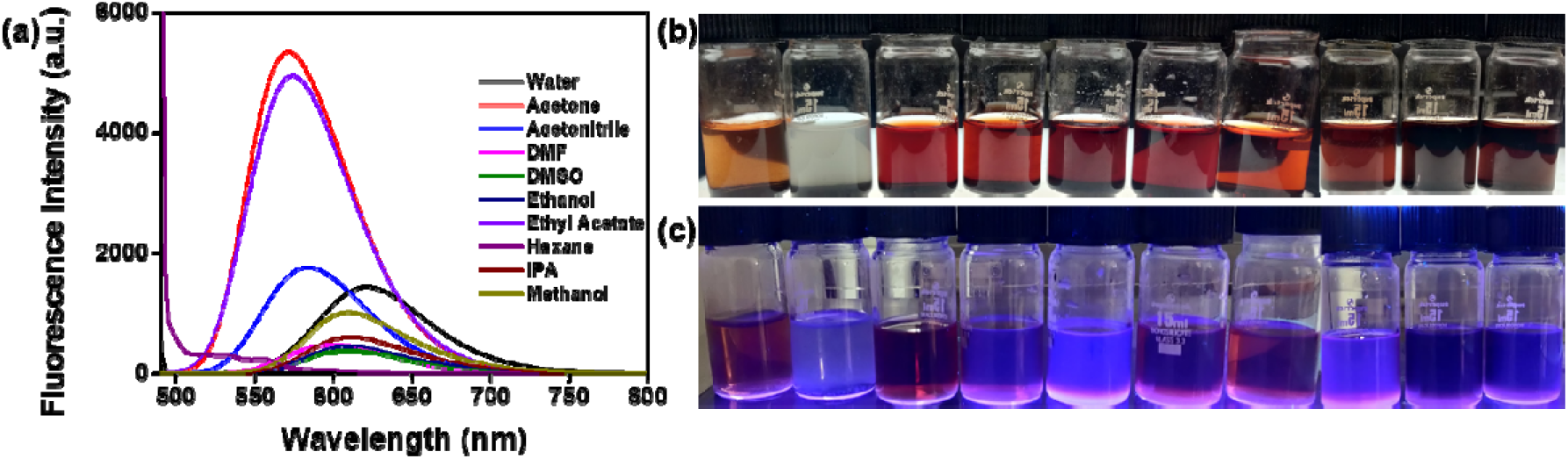
Fluorescence intensity of CNPs in different solvents. **SI 1(a)** Emission spectra of CNPs (0.5 mg/ml) dispersed in different solvents (Water, Acetone, Acetonitrile, DMF, DMSO, Ethanol, Ethyl Acetate, Hexane, Isopropinoic Acid, Methanol) **(b) & (c)** Images of CNPs dissolved in different solvents under white light (above) and UV light (below). (From left to right – Water, Hexane, Acetone, Acetonitrile, DMSO, DMF, Ethyl acetate, IPA, ethanol, methanol)

**SI 2:**
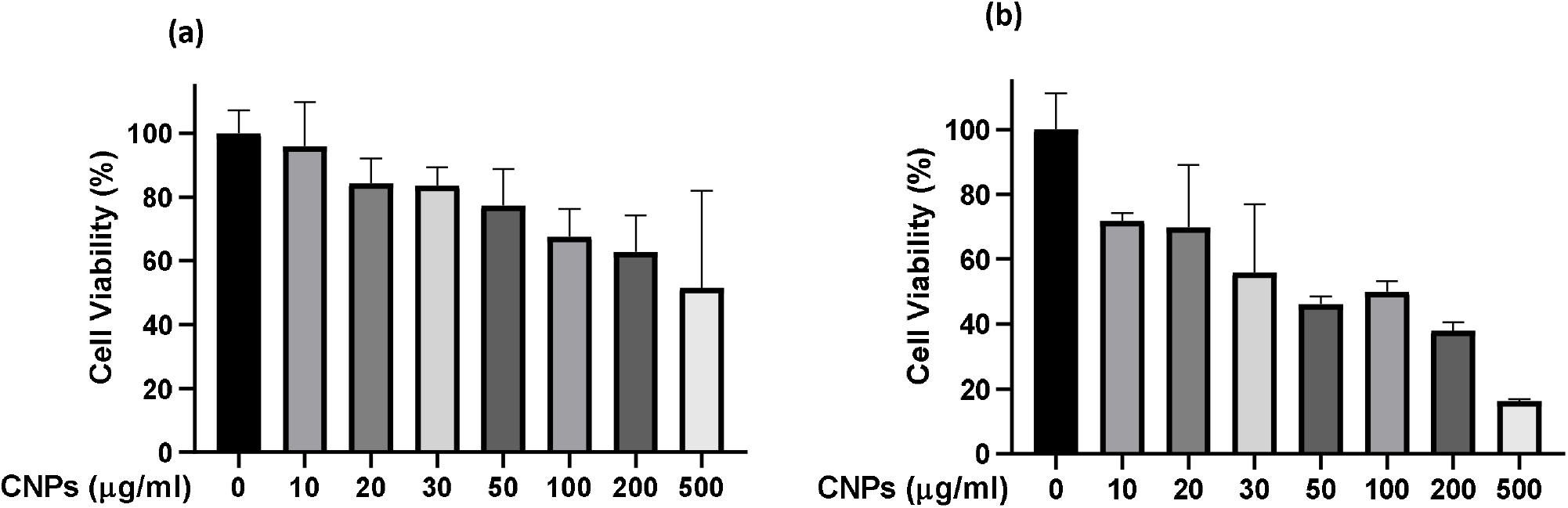
MTT assay. **Figure SI 2**: Cytocompatibility studies. Cell viability was determined by MTT assay using **(a)** SUM 159 and **(b)** MeFs. The cells were treated with CNPs at 10, 20, 30, 50, 100, 200 and 500 µg/ml for 24 h.

**SI 3:**
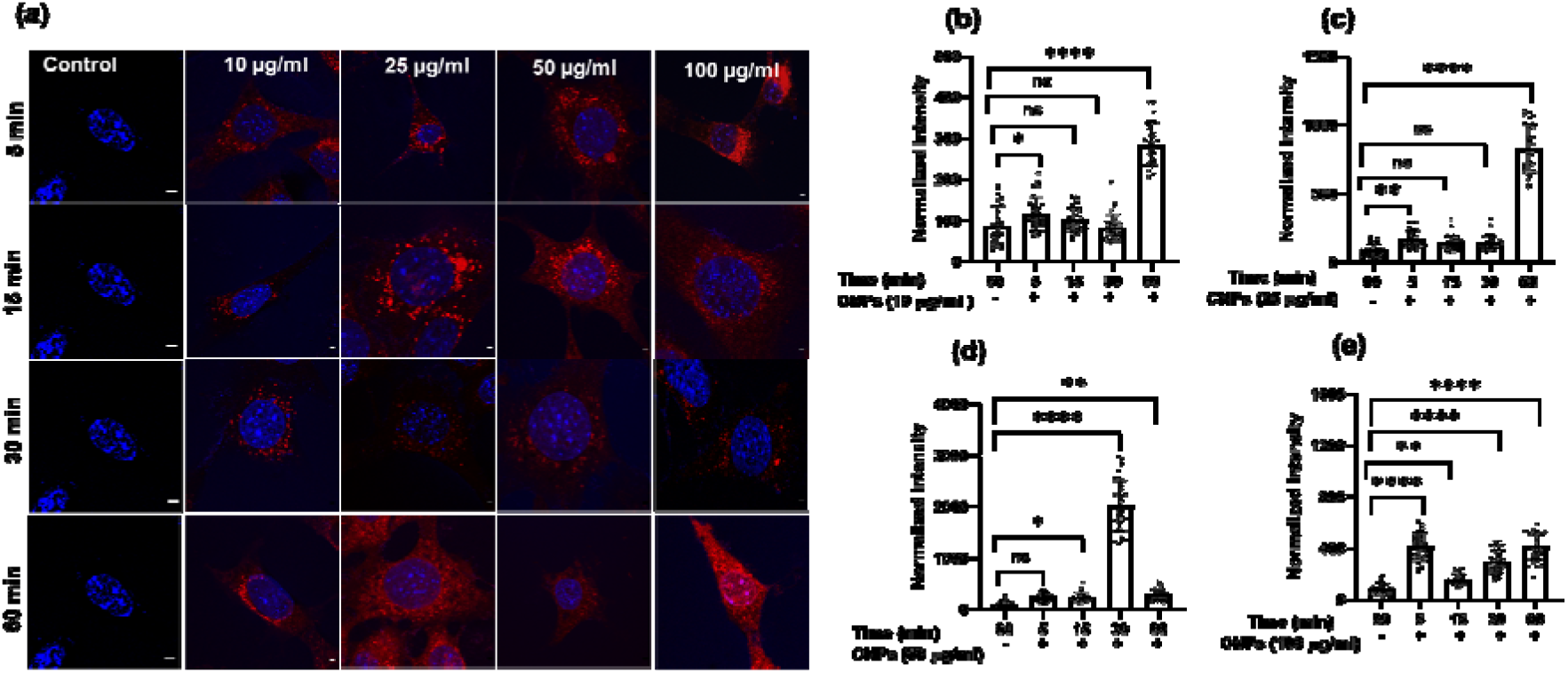
2D Cellular studies of CNPs via confocal microscopy. **Figure SI 3(i):** Concentration dependent cellular uptake studies of CNPs at different time intervals. (a) Confocal images of MeFs incubated with 10, 25, 50 and 100 µg/ml of CNPs respectively. First column in the figure represents untreated MEF cells imaged after 60 min of incubation. Scale bar is 5 µm. (b-e) Quantification of cellular uptake of at 10, 25, 50 and 100 µg/ml of CNPs at 5, 15, 30 and 60 min respectively. **** Indicates statistically significant value of p < 0.0001. ** Indicates statistically significant value of p = 0.002, * Indicates statistically significant value of p = 0.03 and ns indicates non-significant value of p (one-way ordinary ANNOVA).

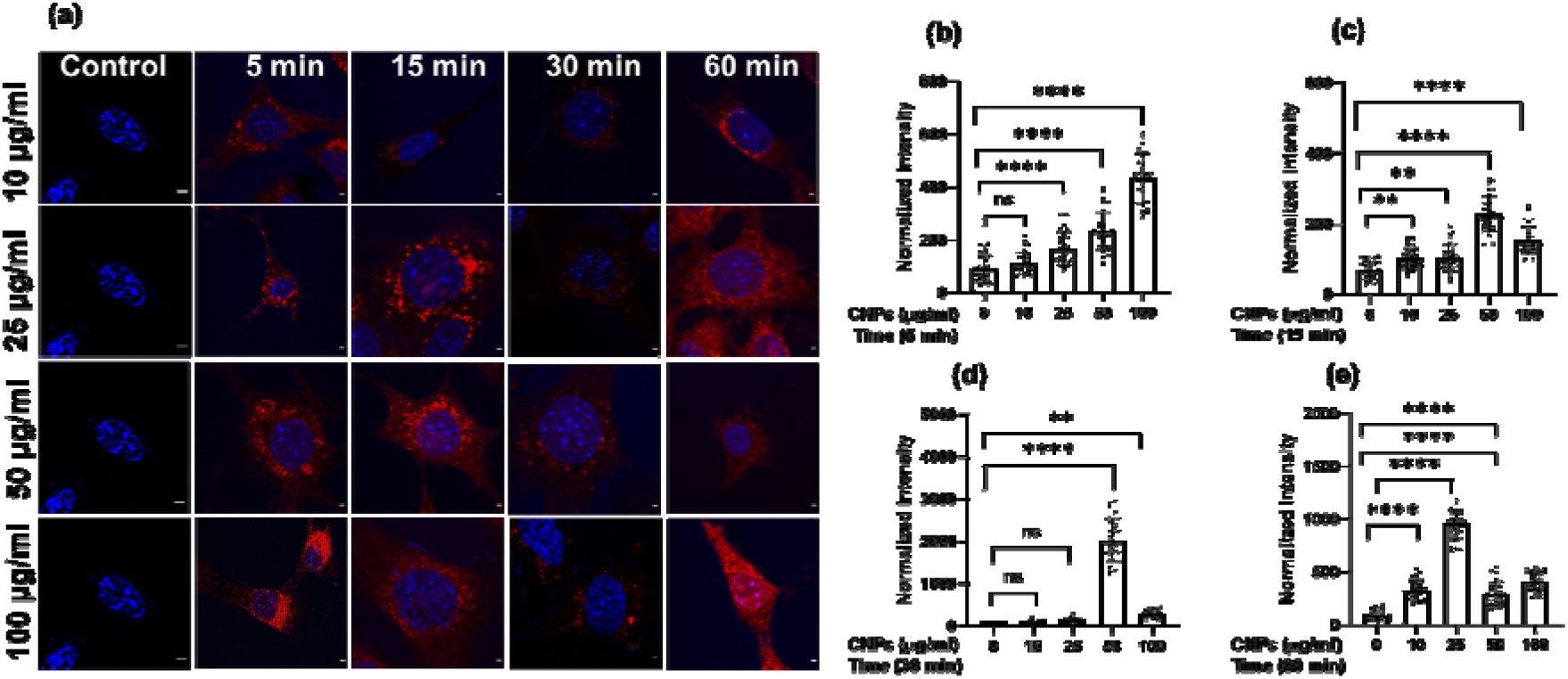
**Figure SI 3(ii):** Time dependent cellular uptake studies of CNPs at different time intervals. (a) Confocal images of MEFs incubated with 10, 25, 50 and 100 µg/ml of CNPs respectively. First column in the figure represents untreated MEF cells imaged after 60 min of incubation. Scale bar is 5 µm. (b-e) Quantification of cellular uptake of CNPs at 10, 25, 50 and 100 µg/ml respectively. **** Indicates statistically significant value of p < 0.0001, ** Indicates statistically significant value of p = 0.005 and ns indicates non-significant value of p (one-way ordinary ANNOVA).

